# Homeostatic plasticity rules control the wiring of axo-axonic synapses at the axon initial segment

**DOI:** 10.1101/453753

**Authors:** Alejandro Pan-Vazquez, Winnie Wefelmeyer, Victoria Gonzalez Sabater, Juan Burrone

## Abstract

GABAergic interneurons are chiefly responsible for controlling the activity of local circuits in the cortex^1,2^. However, the rules that govern the wiring of interneurons are not well understood^3^. Chandelier cells (ChCs) are a type of GABAergic interneuron that control the output of hundreds of neighbouring pyramidal cells through axo-axonic synapses which target the axon initial segment (AIS)^4^. Despite their importance in modulating circuit activity, our knowledge of the development and function of axo-axonic synapses remains elusive. In this study, we investigated the role of activity in the formation and plasticity of ChC synapses. *In vivo* imaging of ChCs during development uncovered a narrow window (P12-P18) over which axons arborized and formed connections. We found that increases in the activity of either pyramidal cells or individual ChCs during this temporal window resulted in a reversible decrease in axo-axonic connections. Voltage imaging of GABAergic transmission at the AIS showed that axo-axonic synapses were depolarising during this period. Identical manipulations of network activity in older mice (P40-P46), when ChC synapses are inhibitory, resulted in an increase in axo-axonic synapses. We propose that the direction of ChC plasticity follows homeostatic rules that depend on the polarity of axo-axonic synapses.

## Main Text

Homeostatic forms of plasticity are thought to play a central role in stabilising the overall activity levels of neuronal circuits in the brain ^5,6^. Although GABAergic interneurons directly modulate ongoing circuit activity through local connections ^7^, the rules that drive the plasticity of GABAergic synapses are not well understood ^3,8,9^. This question is all the more important in the context of brain development, where GABA released from interneurons switches polarity with age, transitioning from excitation to inhibition during circuit wiring ^10^. Understanding the rules that drive interneuron plasticity will not only shed light on how GABAergic synapses modulate local networks, but also on how circuits in the brain remain stable over time.

Chandelier cells (ChCs) comprise a well-defined class of fast-spiking GABAergic interneurons with axonal arbours that are ideally placed to control neuronal and circuit activity in the cortex ^11,12^. Abnormalities in axo-axonic connections between ChCs and pyramidal neurons have been associated with developmental brain disorders such as schizophrenia ^13^ and epilepsy ^14,15^, but the normal wiring mechanisms of these contacts remain unknown. We used a transgenic mouse line (Nkx2.1^CreERT2^) ^16^ crossed with a reporter line (Ai9) to image the development of ChC axons in layer 2/3 of the somatosensory cortex *in vivo* (Fig. 1a). This line labels ChCs sparsely, allowing accurate morphological reconstructions of dendritic and axonal arbours. In line with the late arrival of ChCs to the cortex, *in vivo* imaging of individual ChCs over many days also showed a delayed period of axonal growth that peaked within a narrow window across different cells, from P12 to P18. More surprising was the fact that the axonal arbours of individual ChCs showed a rapid transition in their morphology, generally within two days, from an immature state with few cartridges, to a highly complex arbour with multiple cartridges that span a well-defined cortical domain (Fig. 1b, c). The dendrites, on the other hand, appear to develop earlier and remain largely unchanged throughout this period. Mirroring the rapid growth of the axonal arbour, imaging of fixed brain slices at different developmental periods showed that the number of postsynaptic pyramidal cells contacted by an individual ChC also increased during this window (Fig. 1d, e). Although our data revealed that some connections between ChCs and pyramidal cells existed before P12 (∼50% connectivity on average; Fig. 1e), these connections were generally weak, involving few synapses (Fig. 1f-h). Indeed, we observed an abrupt increase in the number of synapses formed onto an AIS (Fig. 1f-h) that followed the increase in axon arbour size, without any changes in AIS length (Fig. 1i). In agreement with these morphological findings, we also saw a functional increase in the amplitude (Fig. 1j) of GABAergic PSCs recorded from pyramidal cells in response to optogenetic stimulation of ChCs (Extended Data Fig. 1) during the period of synaptogenesis, as well as a maturation of the intrinsic firing properties of ChCs (Extended Data Fig. 2). We conclude that there is a narrow developmental window (P12-P18) over which ChCs connect to neighbouring pyramidal cells and establish a local microcircuit.

**Fig. 1.**
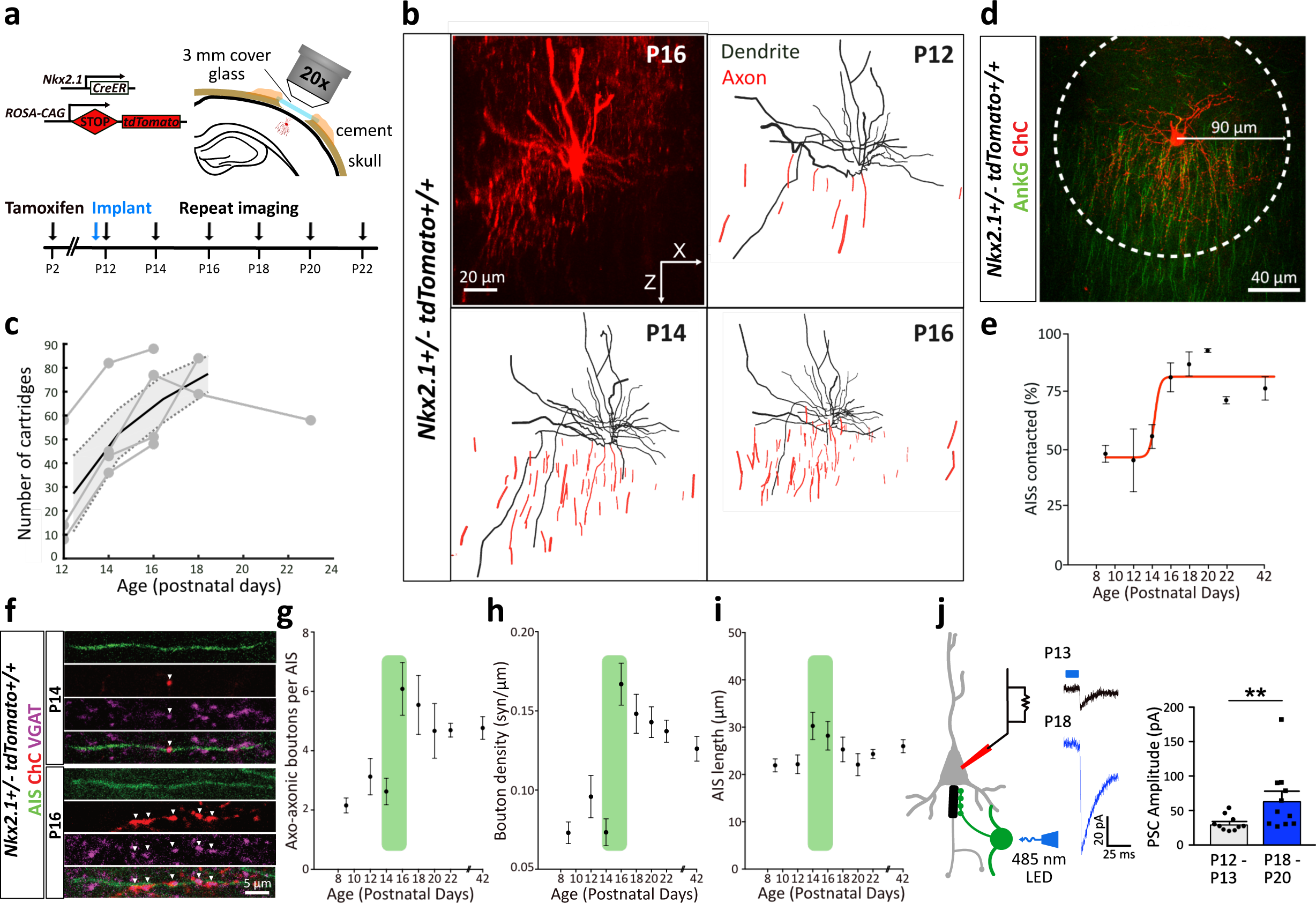
Development of chandelier cells and axo-axonic synapses in somatosensory cortex. **(a)** Genetic strategy and timeline for tamoxifen injection for labelling ChCs in Nkx2.1^+/-^ CreERT2:Ai9 mice, cranial window implantation and repeated *in vivo* imaging. **(b)** *In vivo* image (P16) and reconstructions (P12-P16) of a ChC. **(c)** Number of cartridges for individual ChCs during development (grey) and mean cartridge number (black, n=4 ChCs, 3 mice). **(d)** Image of a ChC (red) and AISs (green) at P18. Connection probability was defined as the percentage of AISs with ChC overlap within a 90 µm radius (white circle). **(e)** Average connection probability of ChCs across development (4-5 ChCs, 2-4 mice per time-point), with a sigmoidal fit (red). **(f)** Images of axo-axonic synapses located on an AIS and expressing VGAT at P14 and P16. **(g**) Average number and **(h)** density of axo-axonic boutons as well as **(i)** AIS length across development, from fixed tissue samples (N=26-81 AISs, 2-4 mice per time-point). The green shaded area highlights the period of rapid synaptic development. **(j)** ChCs expressing ChR2 were stimulated with light and GABAergic PSCs recorded in nearby pyramidal cells (left). Example responses (middle) and average GABAergic PSC amplitude (right) in immature and mature networks (**p<0.01, Mann-Whitney test. N = 10-11 neurons, 4 mice, per condition). Plots show mean ± s.e.m.

We next explored the role that network activity plays in the formation of ChC circuits in the somatosensory cortex. Using designer receptors exclusively activated by designer drugs (DREADDs - specifically hM3Dq) ^17^ expressed in layer 2/3 pyramidal neurons in the somatosensory cortex, we increased network activity during the window of ChC synaptogenesis by delivering the DREADD agonist, clozapine-N-oxide (CNO), from P12 to P18 (Fig. 2a). This manipulation resulted in an increase in activity (verified by cFos expression, Extended Data Fig. 3) in both pyramidal cells that expressed hM3Dq and neighbouring cells that did not (referred to as hM3Dq-network), suggesting a network-wide increase in neuronal activity. Indeed, although increases in activity were initially confined exclusively to DREADD-expressing neurons (at P12; Extended Data Fig. 3a-c), activity spread to neighbouring neurons in the network by the end of the CNO treatment (at P18; Extended Data Fig. 3d, e). We found that increased activity during this period resulted in a decrease in the overall number of pyramidal neurons contacted by a single ChC (Fig. 2b), a finding that highlights the role of activity in moulding the connectivity of inhibitory microcircuits. In parallel to this, we found that the number of axo-axonic synapses formed by single ChCs decreased significantly in both pyramidal neurons expressing DREADDs, as well as in neighbouring DREADD-negative (hM3Dq-network) pyramidal cells (Fig. 2 c-g), in agreement with a network-wide increase in activity (Extended Data Fig. 3d, e). The connectivity between ChCs and pyramidal neurons was further studied functionally. GABAergic PSCs were recorded in pyramidal cells in response to optical stimulation of ChCs expressing channel-rhodopsin-2 (ChR2). Again, matching the morphological decreases in axo-axonic boutons, we found that the amplitude of evoked GABAergic PSCs also decreased, as did the failure rates of synaptic transmission, in line with the idea that few synapses (∼5 boutons per ChC-pyramidal cell contact; Fig. 2e-g) contribute to the postsynaptic response (Fig. 2h-k). Finally, although both morphological (Fig. 2b) and functional (Fig. 2l) measures of connectivity decreased significantly, the absolute values differed in each case, a discrepancy that is likely explained by the unbiased low stringency rules used to measure morphological connectivity (see methods). We conclude that axo-axonic synapses are sensitive to network activity during this early period of synapse formation, decreasing their output in a hyperactive environment. Interestingly, basket cells, fast spiking interneurons that predominantly target the soma, showed an increase in the number of boutons onto hyperactive pyramidal cells (Extended Data Fig. 4), suggesting differences in either the synaptic properties or the plasticity rules between interneuron subtypes during this period. Finally, measures of the structural properties of the AIS showed that DREADD-expressing pyramidal cells (but not neighbouring hM3Dq-network cells) tended to have shorter AISs (Extended Data Fig. 5a), which were matched by a decrease in intrinsic excitability (Extended Data Fig. 5b), a finding that is in line with homeostatic forms of plasticity previously observed at the axon initial segment in other systems ^18-22^. The confinement of AIS plasticity to DREADD-expressing neurons only, is puzzling and suggests that higher levels of activity are needed to drive this form of plasticity, which may only be achieved in those pyramidal cells directly activated by CNO. It also suggests that distinct mechanisms are likely used for AIS versus axo-axonic synapse plasticity ^21^.

**Fig. 2.**
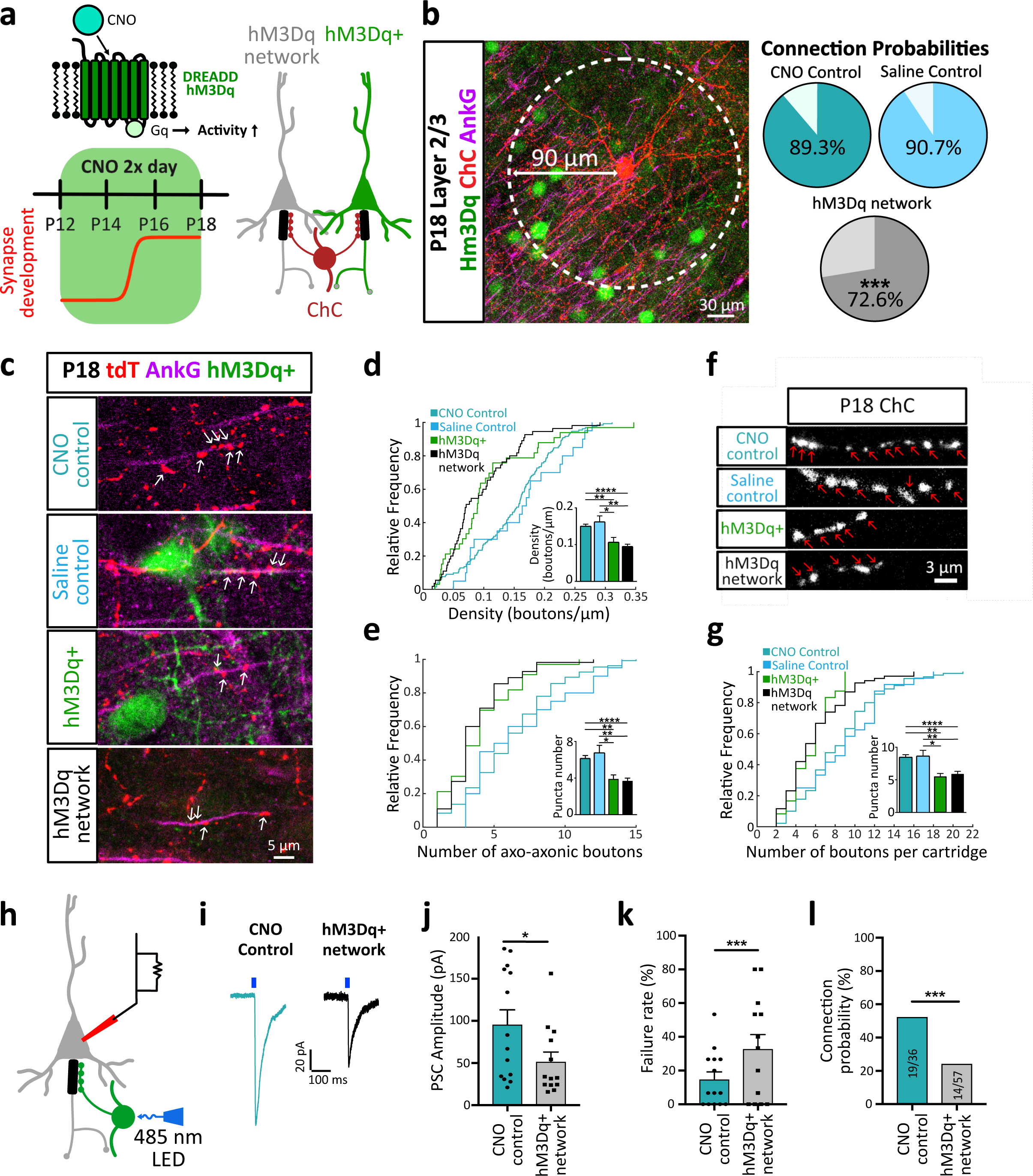
Activity-dependent plasticity of axo-axonic synapses. **(a)** Schematic of DREADD receptor hM3Dq (top left), timeline of CNO application (below) and logic of experimental design resulting in ChCs contacting hM3Dq+ (green) and hM3Dq-(grey) cells in the same network. **(b)** Connection probability of ChCs at P18 in a 90 µm radius (white circle) following injection of CNO into control mice (CNO control), into hM3Dq expressing mice (hM3Dq network) or saline into hM3Dq expressing mice (saline control) (Chi-Square test. N=270-390 cells, 3-4 mice, per condition). **(c)** Example images showing axo-axonic boutons overlapping with the AIS after treatment at P18. **(d-e)** Cumulative distribution of and average axo-axonic bouton density and number (One-way Kruskal-Wallis with Dunn’s multiple comparison test. N=20-132 cells, 3-4 mice, per condition). **(f)** Example images of axo-axonic cartridges. **(g)** Cumulative distribution and average cartridge size (Kruskal-Wallis with Dunn’s multiple comparison test, N=21-40 neurons, 3-4 mice, per condition). **(h,i)** Representative GABAergic PSCs from pyramidal cells after optogenetic ChC stimulation for CNO control (blue) and hM3Dq-network (black) conditions. **(j)** Average GABAergic PSC amplitudes and **(k)** average failure rates (Chi-Square test, N=13-14 neurons, 4 mice, per condition). **(l)** Connection probability (Chi-Square test, N=36-57 neurons, 5 mice, per condition). Bar plots show mean ± s.e.m.

The plasticity of both axo-axonic synapses and of the AIS were found to be reversible. Following six days of increased activity, mice were allowed to recover for a further five days without any CNO injections, after which axo-axonic synapse properties were assessed (Extended Data Fig. 6a). All measures of axo-axonic synapse connectivity were found to recover back to normal levels, indistinguishable from unstimulated neurons of the same age (Extended Data Fig 6b-e). Surprisingly, although the length of the AIS also recovered, it grew beyond control levels (Extended Data Fig. 6f), suggesting some kind of rebound effect during the recovery period. Together, our findings show that network activity reversibly controls the output of pyramidal cells during development by modulating both the structure of the AIS as well as the axo-axonic synapses that form onto it.

Increases in network activity are also likely to drive activity in ChCs through local intracortical connections ^23^. To establish if axo-axonic synapse plasticity requires the activity of pyramidal cells or can be driven by exclusively increasing the activity of ChCs in a cell-autonomous manner, we expressed DREADDs (hM3Dq) solely in ChCs (Fig. 3a, b). The low efficiency of this approach also resulted in very sparse expression patterns, typically labelling a single ChC within the somatosensory cortex. Activation with CNO resulted in a similar reduction in synapse number and connection probability when compared to the plasticity driven by increases in pyramidal cell activity (Fig. 3c-f). This manipulation did not result in any significant changes in the size of the AIS of pyramidal neurons (Extended Data Fig. 7). Our results are consistent with the idea that ChCs sample the ongoing levels of activity in local pyramidal cell circuits through changes in their own activity levels and alter their output accordingly.

**Fig. 3.**
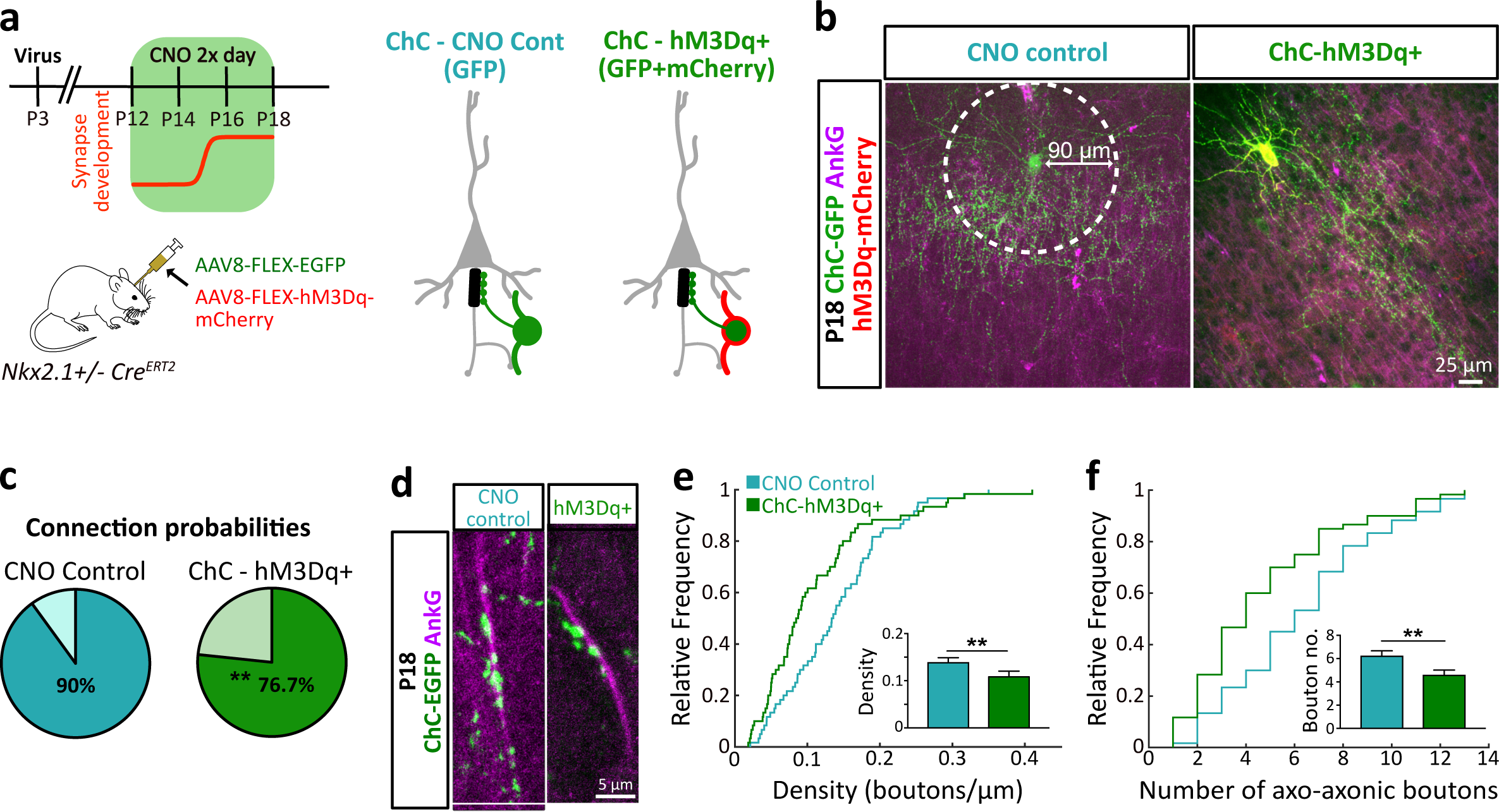
Axo-axonic plasticity is cell-autonomous. **(a)** Timeline of CNO delivery (top left) and strategy for viral delivery of GFP and hM3Dq to ChCs (bottom left). Experimental conditions of viral strategy (right). For CNO control, ChCs were infected with a GFP virus only, whereas for the hM3Dq+ condition, ChCs were infected with GFP and hM3Dq-mCherry. **(b)** Example images of P18 control and hM3Dq+ ChCs. **(c)** Connection probability within a 90 µm radius (white circle in b) for control and hM3Dq+ ChCs. Chi-square test, N=120-150 AISs from 3 mice, per condition. **(d)** Example images of P18 control and hM3Dq+ axo-axonic boutons. **(e,f)** Cumulative distribution of and average axo-axonic bouton density and number. Mann-Whitney test, N = 60 AISs from 3 mice, per condition. Bar plots show mean ± s.e.m.

In the context of homeostatic forms of plasticity that are thought to operate over these long timescales ^5,24^, the decrease of ChC synapses in hyperactive networks during development (Fig. 2a-g) is surprising and in clear contrast to the increase in somatic GABAergic inputs in the same network (Extended Data Fig. 4). This discrepancy may be explained by recent work studying the switch in polarity of GABAergic synapses, as they transition from excitatory to inhibitory/shunting during development, along different subcellular compartments ^25^. Whereas GABAergic synapses on the somato-dendritic compartment switch polarity within the second postnatal week ^10^, those on the AIS remain depolarising well into the third postnatal week, providing a temporal window over which different interneurons could potentially drive pyramidal cell activity in opposite directions ^25^. We therefore explored the polarity of axo-axonic synapses at the AIS during this early period of synaptogenesis by performing voltage imaging of pyramidal neurons in acute slices obtained at P16-P18. This approach provided a readout of membrane potential responses to either local GABA iontophoresis or ChC stimulation without perturbation of the intracellular milieu. We first carried out local GABA iontophoresis at the AIS of pyramidal neurons expressing the genetically-encoded voltage indicator Ace-mNeon ^26^ (Fig. 4a; see Extended Data Fig. 8 for voltage sensitivity). In agreement with previous findings ^25^, GABA tended to produce strong depolarisations at the AIS that were sensitive to GABA_A_ receptor antagonists during this developmental period (Fig. 4b, c; note that depolarisations produce a drop in fluorescence). To explore this further, we performed whole-cell patch clamping of ChCs to directly stimulate a single ChC and image voltage responses in neighbouring pyramidal cells that expressed Ace-mNeon (Fig. 4d). Although GABAergic postsynaptic potential responses were small (Fig. 4e), we saw a clear overall depolarisation above baseline during the stimulation period (5 APs at 50 Hz) that was apparent in individual cells, as well as in the mean response across all 22 cells (Fig. 4f). Our results strongly suggest that release of GABA by ChCs causes postsynaptic depolarisations in pyramidal neurons during the early period of synaptogenesis (P12-P18), in agreement with others ^25,27-29^. These findings have important implications for interpreting the plasticity of axo-axonic synapses. We propose that the decrease in depolarising axo-axonic synapses in response to chronic elevation of activity is a homeostatic response that aims to stabilise circuit activity in the cortex.

**Fig. 4.**
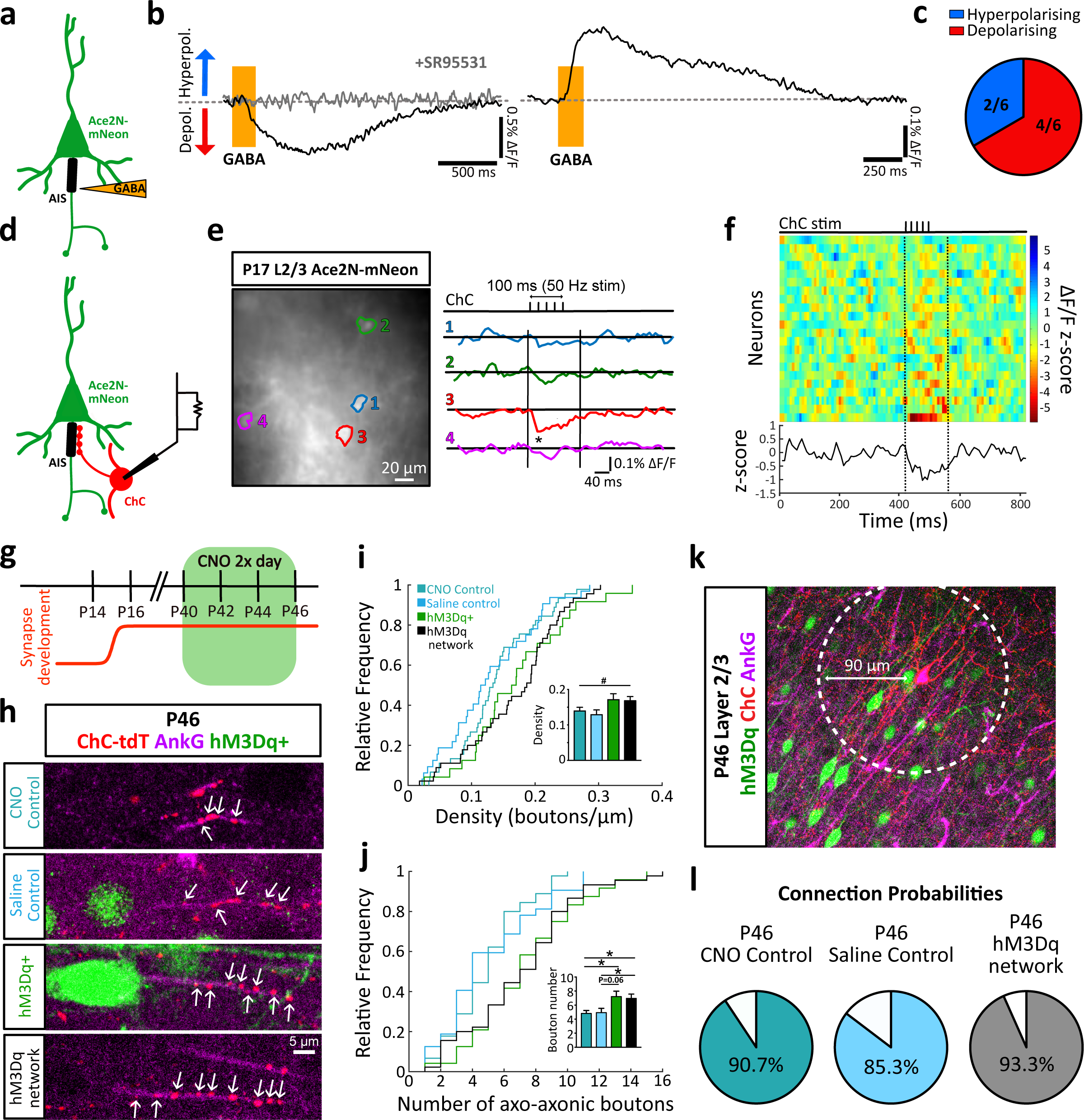
Axo-axonic plasticity matches GABAergic polarity. **(a)** Schematic showing iontophoresis experiments. Pipettes containing GABA were placed near the AIS of a pyramidal neuron expressing Ace2N-mNeon. **(b)** Example traces obtained from imaging somatic responses to iontophoretic GABA application (orange area) at the AIS of two different cells. Left, a depolarising response that is blocked by the application of 10 µM SR95531 (GABAzine). Right, example trace showing a hyperpolarising response. **(c)** Classification of responses into hyperpolarising (blue) and depolarising (red) for all cells tested. **(d)** Schematic showing experimental logic: ChCs were patch-clamped to evoke APs and responses were imaged from the soma of nearby pyramidal cells expressing Ace2N-mNeon. **(e)** Left, example image of Ace2N-mNeon cells, with somas outlined in different colours. Right, change in fluorescence of corresponding cells following ChC stimulation (top; 5 APs, 50 Hz) (* denotes time-locked event 3x > baseline standard deviation). Vertical lines denote start and end of expected response period, allowing up to 50 ms for decay of GABAergic responses (Fig. 2i) **(f)** Heat map showing z-scores of fluorescence responses following ChC stimulation for all 22 pyramidal cells tested. Average z-score across all cells is shown below. **(g)** Timeline for CNO delivery in adult mice. **(h)** Example images of axo-axonic boutons in the different conditions at P46. **(i,j)** Cumulative distribution and average axo-axonic bouton density and number (# denotes p<0.05 for one-way ANOVA without reaching significance in posthoc comparisons; *p<0.05 denotes Kruskal-Wallis and Dunn’s posthoc comparisons test, N=44-72 neurons per condition, 3-5 mice, per condition). **(k,l)** Connection probability within a 90 µm radius (white circle) at P46 (Chi-square test, N=150-180 AISs, 3-4 mice, per condition). Bar plots show mean ± s.e.m.

Previous studies have shown that ChCs in adult brains are inhibitory, rather than excitatory^23,25^. We therefore reasoned that if ChCs follow homeostatic rules to control axo-axonic synapse number, the same increase in network activity performed in older animals should result in the opposite phenotype to that observed during development. Indeed, increasing the activity of cortical networks by delivery of CNO to DREADD (hM3Dq)-expressing pyramidal neurons from P40 to P46 resulted in an increase in axo-axonic synapses along the AIS (Fig. 4g-j). Unlike manipulations during early development, the plasticity of axo-axonic synapses was not accompanied by changes in the size of the AIS (Extended Data Fig. 9) nor in the number of postsynaptic pyramidal cells contacted by a single ChC (Fig. 4k, l). The switch in the plasticity of ChC outputs can be explained as a homeostatic response that depends on the polarity of the synapse at the time. Whereas in mature brains axo-axonic synapses are inhibitory and therefore respond to hyperactivity by increasing their number, early in development, when they are depolarising, they respond by decreasing their drive onto pyramidal cells. The direction of synaptic plasticity is therefore dependent on the polarity of axo-axonic synapses. Similarly, the opposite direction in the plasticity of axo-axonic versus axo-somatic synapses early in development is likely driven by the difference in polarity of each synapse type at that developmental stage.

Our results suggest that the wiring of GABAergic interneurons in the cortex is governed by homeostatic rules that control the number and strength of connections onto pyramidal cells. In this context, the plasticity of the AIS, together with its axo-axonic synapses, provides a hub for tightly modulating pyramidal cell activity, where multiple forms of plasticity come together in a concerted manner to stabilise the output of pyramidal neurons in the cortex. The overall functional consequence of this arrangement underscores the importance of GABAergic interneurons in maintaining network stability during the highly dynamic period of circuit formation.

## Acknowledgments

We would like to thank Matthew Grubb, Adil Kahn, Oscar Marin and Beatriz Rico for comments on the manuscript; Ruben Deogracias and Beatriz Rico for help with the virus injections; Varun Sreenivasan and Oscar Marin for help with 2-photon microscopy. This work was supported by a Wellcome Trust Investigator Award (095589/Z/11/Z) and an ERC Starter Grant (282047) to JB, as well as MRC and KCL studentships to APV and to VGS, respectively.

## Contributions

Conceptualization, APV, WW and JB; Methodology, APV, WW, VGS; Software, APV, WW and VGS; Formal analysis, APV, WW and VGS; Investigation, APV, WW; Writing-Original Draft, APV, WW and JB; Writing-Review & Editing, APV, WW and JB; Funding Acquisition, APV, VGS, JB; Supervision WW and JB.

## Materials and Methods

### Animals

Male and female mice were used for all experiments. Mice were housed in individually ventilated cages and provided with *ad libitum* food and water. Delivery of viral vectors was done in the Nkx2.1^+/-^*CreER* line (JAX014552). For morphological analysis and voltage imaging, Nkx2.1^+/-^*CreER* mice were crossed with the Ai9 (JAX007909) line. For measuring the intrinsic excitability of chandelier cells, Nkx2.1^+/-^-*CreER* mice were crossed with both the Ai9 and the Ai32 line (JAX012569). For measuring the electrophysiological properties of the axo-axonic synapses Nkx2.1^+/-^*CreER*/Ai32 mice were used. For measuring spontaneous GABAergic PSC frequency, Nkx2.1^+/-^-*CreER*/Ai32 were used. Finally, for recording the intrinsic excitability of pyramidal cells after DREADDs treatment and iontophoresis experiments, Ai9 mice were used. Both homozygous and heterozygous mice were used. All procedures were carried out in accordance with the UK Animal Scientific Procedures Act.

### Tamoxifen induction

For the optimal labelling of ChCs, Tamoxifen dependent Cre recombination was induced at P2. For experiments using *Cre*-dependent viral vectors, Tamoxifen was injected at P3, on the day of viral injection. Tamoxifen (Sigma) was dissolved in corn oil (30 mg/mL) and injected intraperitoneally at a dose of 100 µg/g.

### Cranial windows

Cranial windows were implanted covering somatosensory cortex as in previous studies *(28)*. P11-P12 mice were anaesthetised with 2% isoflurane in a nose-clamp and body temperature maintained at 37°C with a heating pad. The local anaesthetic Bupivacaine was injected sub-cutaneously (10 µl of 0.5 µg/ml) and the anti-inflammatory dexamethasone was applied intramuscularly (10 µl of 38 µg/ml solution) at the start of the procedure. The scalp was cleaned with betadine and cut open to expose the cranium. Lidocaine Hydrochloride (1% w/v) was then applied briefly to the cranial surface before the periosteal tissue was carefully removed with a scalpel. The cranial surface was then cleaned with Ringer’s solution and a few drops of betadine. A custom-made metal head-post (design by Carl Petersen lab) was glued to the right hemisphere with cyanoacrylate glue (Henkel). Dental cement (Paladur) was applied around the head-post and the cranium to reinforce the implant. A 3 mm craniotomy was opened over the somatosensory cortex with a dental drill until the dura was exposed. Care was taken not to cross the bone sutures. Cortex buffer (NaCl [123.35 mM], KCl [5 mM], Glucose [10 mM], HEPES [10 mM], CaCl2 [2 mM], MgSO4 [2 mM]) was applied to the exposed surface and a 3 mm glass coverslip (Harvard Apparatus) placed directly over the craniotomy, attached at the edges with cyanoacrylate glue and strengthened with dental cement. At the end of the procedure, mice were injected with saline and left to recover for 4 h - 1 day before imaging.

### 2-photon *in vivo* imaging

Mice were anaesthetised with Ketamine/Xylazine. Imaging sessions lasted 2-3 hours. The same field of view was imaged over consecutive days. tdTomato was excited with a Ti-Sapphire laser (Coherent Chameleon) at 930 nm wavelength. Emitted light was collected by a GaAsP detector through a 1.0 NA, 20× objective (Olympus). The excitation power was between 40 and 50 mW.

### Histology

Mice were anaesthetised with an overdose of sodium pentobarbital and transcardially perfused with 10 ml of ice-cold saline solution followed by 20 ml of 1% (w/v) PFA in HEPES (pH 7.3). The brains were carefully removed and put in 1% (w/v) PFA solution overnight at 4°C. Brains were embedded in blocks of 6% agarose (Sigma) and sectioned coronally (100 µm thick) with a vibratome (Leica). Slices were kept in PBS supplemented with 0.05% Sodium Azide to improve preservation. On the day of staining, slices were incubated at RT for 2h in PBS-Triton (0.25%) supplemented with 10% Goat Serum (GS) (Sigma). This was followed by an overnight incubation in a primary antibody solution in PBS-Triton (0.25%)-GS (10%). On the following morning, slices underwent three 20 minute PBS-Triton (0.25%) washes, followed by a 2 hour incubation at room temperature in secondary antibody solution in PBS-Triton (0.25%)-GS (10%). The secondary antibody solution was washed off as before and slices were mounted with Vectashield mounting media with DAPI (VectorLabs) and kept at 4°C. The following primary antibodies were used: Anti-AnkyrinG-mouse2b (NeuroMab), Anti-DsRed-Rabbit (Takara), Anti-GFP-Chicken (Abcam), Anti-VGAT-mouse3 (Synaptic Systems), Anti-cFos-mouse1 (Santa Cruz), Anti-NeuN-mouse1 (Millipore). Alexa Fluor conjugated antibodies (Invitrogen) were used as secondary antibodies.

### In utero electroporation

CAG:hM3Dq:IRES:GFP, CAG:hM3Dq:IRES:RFP or CamKII-ACE2N-mNeon-4AA were expressed in pyramidal cells via *in utero* electroporation. *In utero* electroporation was performed in E15.5 pregnant mice. Females were anaesthetised with isoflurane 2.5% before the abdomen was opened and the uterine horns exposed. DNA was delivered to the embryos via injection into the lateral ventricles with a glass micropipette. A total of 0.5-1 µL of DNA solution was injected into each embryo. The DNA solution contained Fast Green (0.3% (w/v)) and DNA in TE buffer (1500-2000 ng/ml). Five square electric pulses (30 V, 50 ms) were passed at 1s intervals using a squarewave electroporator (CUY21EDIT; NEPA GENE).

### Expression of viral constructs in ChCs

AAV8-CAG-FLEX-GFP (UPenn Vector core) and AAV8-hsyn-hM3D(Gq)-mCherry (UNC Vector Core) *(29)* were injected into L2/3 somatosensory cortex of P3 Nkx2.1^+/-^*CreER* mice. Mice were anaesthetised with 2.5% isoflurane. The following coordinates were used: Anterior-Posterior-axis: 2.7 mm, Dorso-Ventral: −0.3 mm, Lateral: −2.3 mm from Bregma. A total volume of 250 nl was infused at a flow rate of 50 nl/min. All injections were performed with a Nanoliter injector (WPI) controlled with a SMARTouch controller (WPI) as in *(30,31)*.

### Clozapine-N-Oxide (CNO) treament

For DREADDS experiments CNO (Enzo Life Sciences) was dissolved in 0.5% dimethyl sulfoxide (DMSO) and 0.9% saline to a final concentration of 0.5 mM. Once in solution, CNO was aliquoted and stored at −20°C until the day of use. Animals were injected intraperitoneally twice daily with saline (0.9%) with DMSO (0.5%) or CNO in DMSO (0.5%) at 1 mg/kg. The saline volume injected was calculated based on the CNO equivalent for the given weight of the animal. On the final day of the experiment animals were only injected once, 2h before perfusion.

### Image acquisition of histology

A Nikon A1R inverted confocal microscope with a 40x water immersion objective (NA 1.1) was used for the acquisition of images from immunostained slices at 1024 × 1024 pixels resolution. For synaptic measurements, a 3x zoom was used. Stacks had a z-step of 1 µm. All images were acquired with NIS Elements software (Nikon).

### Acute slice preparation for electrophysiological and optical recordings

Mice were anaesthetised with Isoflurane and their brains rapidly removed in cold dissection media (Sucrose 240 mM, KCl 5mM, Na_2_HPO_4_ 1.25 mM, MgSO_4_ 2 mM, CaCl_2_ 1 mM, NaHCO_3_ 26 mM and D-glucose 10 mM). 300 µm thick coronal slices were made with a V1000S vibratome (Leica) in constantly carbonated dissection media (95% O_2_, 5% CO_2_). Slices were transferred to an immersion storage chamber containing artificial cerebrospinal fluid (ACSF) with the following composition: NaCl (124 mM), KCl (5 mM), Na_2_HPO_4_ (1.25 mM), MgSO_4_ (2 mM), CaCl_2_ (2 mM), NaHCO_3_ (26 mM), D-glucose (20 mM). Slices were left to recover for 1 hour at room temperature before recording.

### Electrophysiology

Targeted whole-cell patch clamp recordings were made at the soma of ChCs or pyramidal cells in layer 2/3 under voltage and current clamp configurations. Acute slices were transferred to a recording chamber equipped with a BX51 (Olympus) microscope with Dodt gradient contrast optics. For targeted patching, tdTomato+ ChCs were illuminated with a 565 nm LED (2-pE system, CoolLED) which resolved the soma and axonal arbour of the cell. Slices in the recording chamber were constantly superfused with ACSF and carbonated (95% O_2_, 5% CO_2_). For all recordings, the AMPA receptor blocker NBQX (10 µM), and the NMDA receptor blocker APV (25 µM) were present in the bath. Patching electrodes were made from thick-walled borosilicate glass (O.D: 1.5mm, I.D: 0.86 um, Sutter Instruments) and polished with a Microforge MF-900 (Narishige) obtaining a resistance between 3-4 MΩ. Recordings were obtained with a Multiclamp 700B amplifier (Molecular Devices) and digitised with the Digidata 1440A digitizer (Molecular Devices). Data was acquired with the software Clampex (Molecular Devices) and Axon Multiclamp Commander Software (Molecular Devices). Signals were filtered at 10 kHz and sampled at 50 kHz. Pipette offsets were nulled before seal formation and pipette capacitance was compensated in the cell-attached configuration once a giga-seal was obtained. All measurements in voltage-clamp were made after series resistance compensation for 7 MΩ. Recordings were excluded if access resistance exceeded 30 MΩ. All recordings were done at room temperature. Membrane resting potentials were checked in *I* = 0 mode and passive features were recorded on-line. For intrinsic excitability measurements, 10 ms and 500 ms current injections of increasing amplitude (10 pA steps) were injected in current clamp configuration. For measuring GABAergic PSCs, cells were held in voltage-clamp at - 70 mV while ChCs were stimulated with light with 2 ms pulses and 20% LED intensity (470 nm). A 10 s interval was left between trials to allow for the full replenishment of the vesicular pool. For measuring spontaneous GABAergic PSCs, cells were held at −70 mV and recorded for 3 min.

### Internal solutions for patch-clamp recordings

For intrinsic excitability measurements of chandelier and pyramidal cells the internal solution in the recording pipette contained: K-gluconate (130 mM), KCl (15 mM), 10 Na_2_Phosphocreatine (10 mM), HEPES (10 mM), MgATP (4 mM), GTP (0.3 mM) and EGTA (0.3 mM), pH was adjusted to 7.3 with KOH and to 320 mOsm. For recording GABAergic PSCs, a high chloride intracellular solution was used for data in Fig 1 and 2, containing KCl (150 mM), MgCl_2_ (4.6 mM), CaCl_2_ (0.1 mM), HEPES (10 mM), NaATP (4mM), NaGTP (0.4 mM) and EGTA (1 mM), pH was adjusted to 7.3 with KOH and to 320 mOsm. Because of the prevalent presence of action currents in >P18 and older animals, QX-314 (Tocris, 5 mM) was included in the solution for GABAergic PSC recordings in the experiments in Fig 2.

### Voltage imaging

Acute slices were prepared from mice electroporated with the voltage indicator Ace2N-mNeon-4AA in prefrontal and somatosensory cortex. Slices were transferred to a recording chamber equipped with a BX51 (Olympus) microscope with Dodt gradient contrast optics, 470 nm & 565 nm LEDs (CoolLED) and an Evolve 512 EMCCD camera (Photometrics). The slices were continuously superfused with carbonated ACSF containing NBQX (10 µM, Tocris), and APV (25 µM, Tocris). In the iontophoresis experiments the GABA_B_R inhibitor CGP55845 (10 µM, Tocris) was also always present in the bath. Cells were illuminated with a 470 nm LED at 29 mW/mm^2^ (illumination intensity measured at the specimen plane), while imaging at *circa* 120 Hz with an EMCCD camera through a 40x, 0.8 NA water-immersion objective (Olympus). To image at this acquisition rate, 4×4 binning was used, giving a final resolution of 128×128 pixels. Images were acquired using µmanager software (Micro-manager, Vale Lab). Each recording consists of 40-50 repeats with the same stimulus. To obtain a baseline of camera noise, acquisition started 50 ms prior to LED illumination and finished 100 ms after. LED illumination, camera acquisition and stimulation were synchronised with a Digidata 1440A digitizer (Molecular Devices) through the software Clampex (Molecular Devices).

### GABA Iontophoresis

After selecting an ACE-mNeon cell, an iontophoresis pipette was placed juxtaposed to the cell’s AIS or soma. Iontophoresis pipettes were made from thick-walled borosilicate glass (O.D: 1.5mm, I.D: 0.86 um, Sutter Instruments) obtaining a resistance of 60-100 MΩ. The pipettes were filled with a GABA solution containing: GABA (200 mM, pH 5.0), Alexa Fluor 568 (10 µM) and CGP55845 (10 µM, Tocris). Alexa 568 was used to visualise GABA application. CGP55845 was also included in the iontophoretic solution to assure GABA_B_R inhibition from potential displacement of the blocker when GABA was ejected. A retention current of −30 nA was used when the pipette was in the bath to avoid spillage. A single pulse of 20 ms and 150 nA was used to eject the GABA solution from the pipette.

### Morphological analysis of AIS and axo-axonic boutons

For all morphological analyses, care was taken to include an equal number of AISs or cartridges from each animal. AISs and boutons were traced in 3D using the ROI manager and Simple Neurite Tracer (ImageJ). Custom MATLAB code was used to analyse the position of boutons along the AIS and their number. The fluorescent signal for the AIS fades on its ends and has its brightest points at the centre. To normalize AIS length measurements, custom MATLAB code was designed, where the points delimiting the region where intensity dropped to 20% of maximum fluorescence, were assigned as the start and end points. All analyses were performed under blinded conditions. Due to the sparsity of chandelier cells marked in somatosensory cortex, individual cartridges could be related to specific chandelier cells. All chandelier cell varicosities overlapping an AIS were included in the analysis of bouton number and density. To establish a measurement independent of AIS structure, we also measured cartridge size. A cartridge was defined as a collection of 2 boutons or more that project in the same axes as the AIS they synapse onto. This measurement was based purely on axo-axonic cartridge morphology without taking into consideration whether the boutons in the cartridge were contacting an axon. For the analysis of contacts made by an individual chandelier, contact was defined as the presence of at least one ChC varicosity overlapping an AIS. This morphological estimate of connectivity is higher than our electrophysiological estimate, presumably due to this low-stringency approach of considering any morphological overlap a contact. On the other hand, our electrophysiological approach is likely underestimating connectivity due to the combination of a small number of synaptic contacts with a low release probability. It is likely that a combination of the two (overestimation of morphological connectivity and underestimation of functional connectivity) explains the absolute difference in connection probability measured in each case.

### Morphological analysis of basket cell boutons

The number of synapses at an individual soma was analysed in a single plane as in *(32)*. Somas were stained with NeuN and GABAergic synapses with VGAT. Thresholding for VGAT and NeuN immunostaining was adjusted in each individual image to ensure optimal segmentation. Segmentation and synapse counting was done automatically. Synapses were segmented with a watershed algorithm. Synapses were included if they were within a 1 µm radius of the segmented soma. Density of synapses was calculated by dividing synapse number over the perimeter of the soma. All analysis was done under blind conditions.

### Analysis of c-fos immunofluorescence

To analyse the effect of DREADDs and its ligand on activity, mice were injected with CNO or saline 2h before perfusion and stained for c-fos, a gene with activity dependent expression. A fluorescence threshold of 1.2x background fluorescence was set to classify cells as c-fos+. For measuring the effects of CNO on hM3Dq+ cells in hM3Dq-networks, cells were blindly selected before measuring their c-fos immunofluorescence profile.

### Analysis of electrophysiological measurements

Input-output curves were generated with custom MATLAB scripts. Two types were used: 1) output in action potential (AP) frequency as a function of the current injected (pA) 2) output in AP frequency as function of current density (pA/Pf). This is the same as 1) but normalised for the capacitance of the cell, which correlates with its size. For the measurements of the voltage threshold, voltage threshold was determined by time of the first peak of the jerk (third derivative of voltage with respect to time). For the measurement of GABAergic PSC amplitudes, all analysis was done with the IGOR (Wavemetrics) package Neuromatic (33). GABAergic PSCs were averaged over 15 sweeps and the maximum amplitude recorded. For the measurement of spontaneous GABAergic PSCs, template matching to a second exponential curve was used to automatically record the number of events. The parameters for template matching were as follows: rise time, 1 ms, decay time, 20 ms, zero-baseline before waveform, 1 ms, total template waveform time, 40 ms. For the measurement of release failures, only cells with at least two successfully evoked postsynaptic potentials were included. Failures were defined as the absence of postsynaptic potentials in response to light stimulation on the presynaptic ChC.

### Analysis of voltage imaging recordings

For analysing voltage imaging recordings from evoked ChC activity, an ROI was selected covering the soma of the pyramidal cell of interest. From this ROI, the fluorescence intensity profile over time for each repeat (stimulation protocol repeat) was obtained. Camera noise was estimated as the average over a 10 frame window with no LED illumination and was subtracted from the profile. The resulting traces were parsed through a custom-made bleach-correction algorithm (MATLAB). In this algorithm, the intensity profile for each repeat was fitted to a double exponential curve excluding the points corresponding to expected evoked response time. From the resulting trace ΔF/F was calculated. F was determined as the average bleach-corrected baseline fluorescence of the last 10 frames prior to the evoked event. The resulting ΔF/F traces for each repeat were then averaged. Finally, the averaged trace was smoothed by applying a 3-frame moving average filter. For analysing voltage imaging recordings from the iontophoresis experiments, a slight variation to this algorithm was introduced to enable bleach correction for traces with long responses and little baseline for fitting. In these experiments the bleach correction algorithm was trained in non-stimulation repeats in the same cells, where no GABA was released.

### Statistics

Statistical analysis was performed with Prism 6 (GraphPad), MATLAB and custom python scripts using the scipy.stats package. Before hypothesis testing, D’Agostino normality test was used in all data sets to determine whether the data followed a normal distribution. Depending on the outcome, the parametric One-Way-ANOVA or non-parametric Kruskal-Wallis test was used when comparing over more than 2 groups. This was followed by a multiple comparison post-hoc analysis with a multiple comparisons correction. For comparison between two groups, a Student’s t-test or the non-parametric Mann-Whitney test was used, depending on normality. Correlations were tested using Pearson’s or Spearman’s method depending on normality. For comparisons of frequency, a X^2^ test was used. Values are presented as mean ± s.e.m. Statistical significance was established at p<0.05.

**Extended Data Figure 1.**
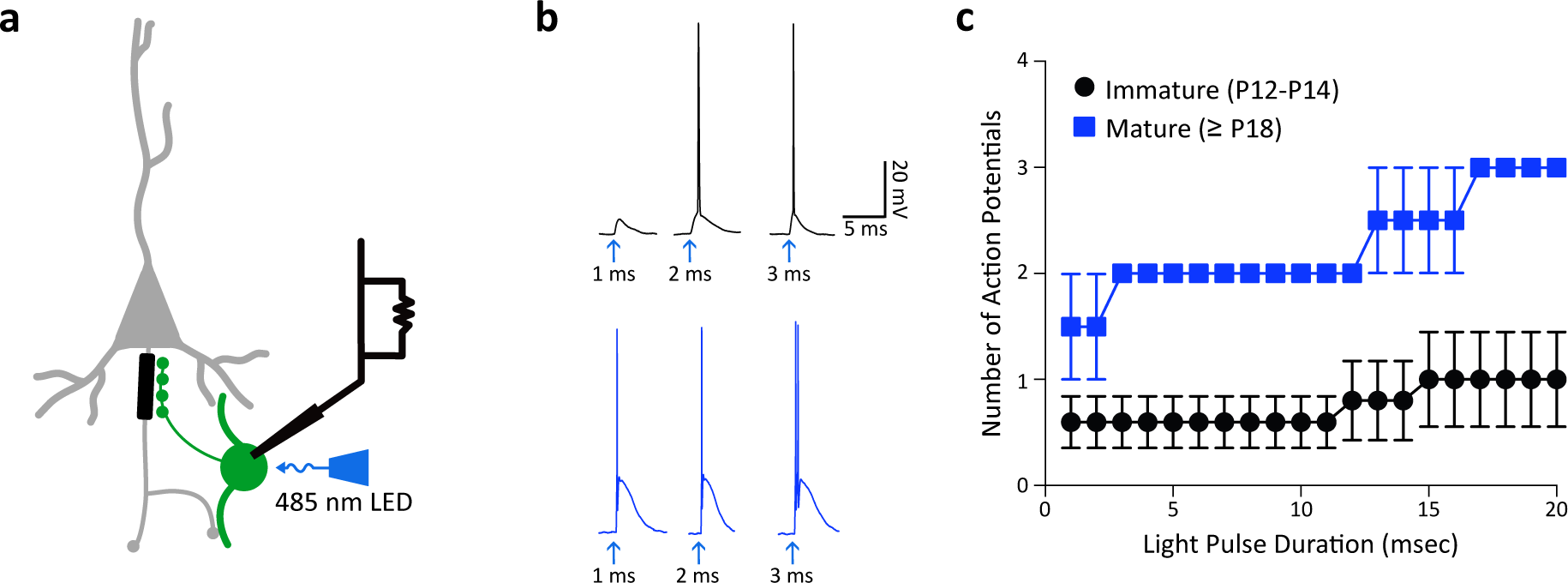
Optogenetic stimulation of ChCs. **(a)** Chandelier cells were patched in whole-cell configuration from Nkx2.1^+/-^CreERT2:Ai32 mice and stimulated with a 470 nm LED. **(b)** Representative action potentials generated with 1, 2 and 3 ms light pulses (arrows) in immature (black) and mature (blue) ChCs. **(c)** Input-output curves showing action potential number as a function of light pulse length in immature (black) and mature (blue) ChCs. Values shown are mean ± s.e.m. Immature: n = 5 ChCs, n = 3 mice; Mature: n = 2 ChCs, n = 2 mice.

**Extended Data Figure 2.**
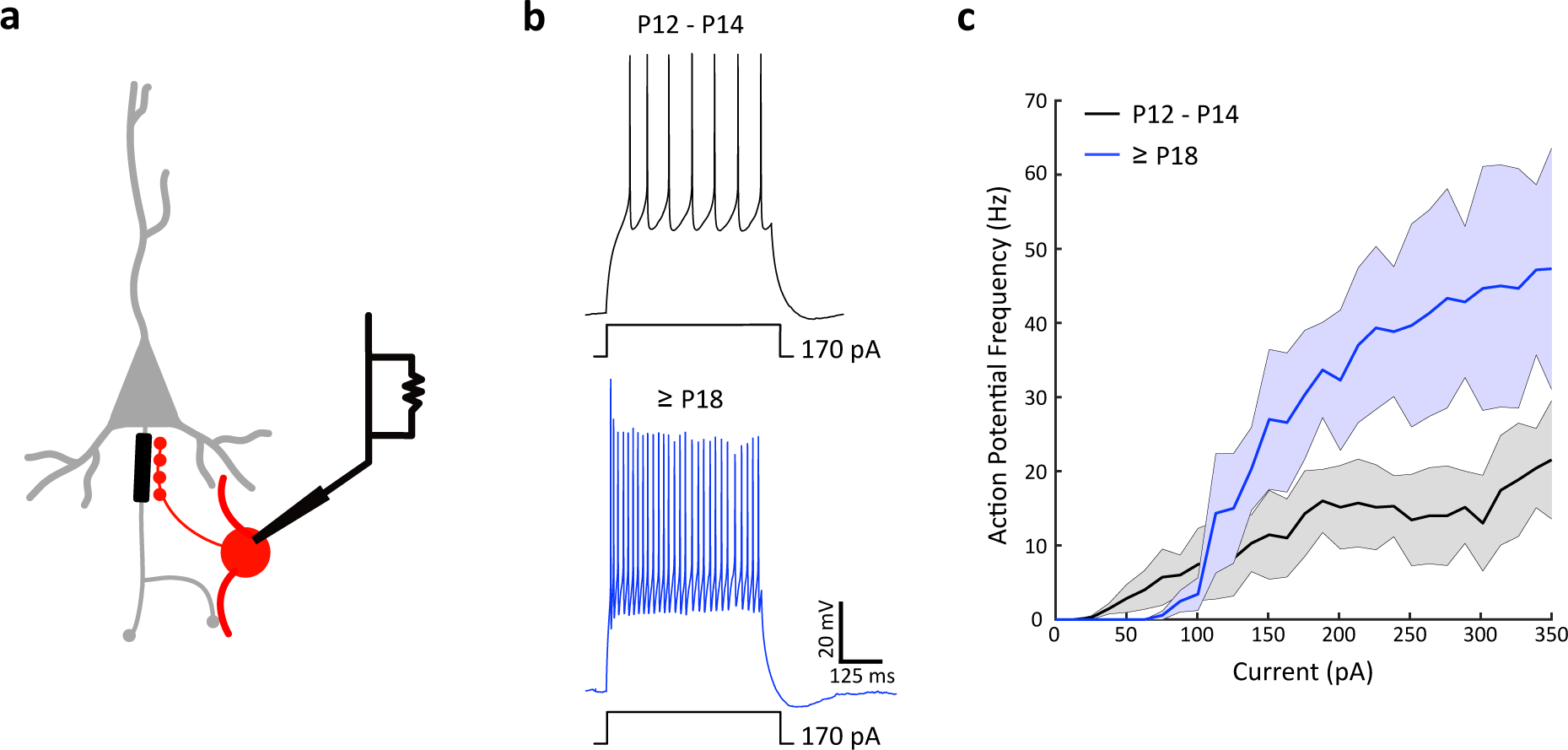
Excitability of ChCs throughout development. **(a)** Chandelier cells were patched in whole-cell configuration during development. **(b)** Representative traces from 500 ms current injections in immature (P12-P14, black) and mature (≥P18, blue) ChCs. **(c)** Input-output curves showing action potential frequency in response to the current injected. Centre line for each group represents mean firing rate, shadow represent s.e.m. Immature: n = 7 ChCs, n = 3 mice; Mature: n = 6 ChCs, n = 4 mice.

**Extended Data Figure 3.**
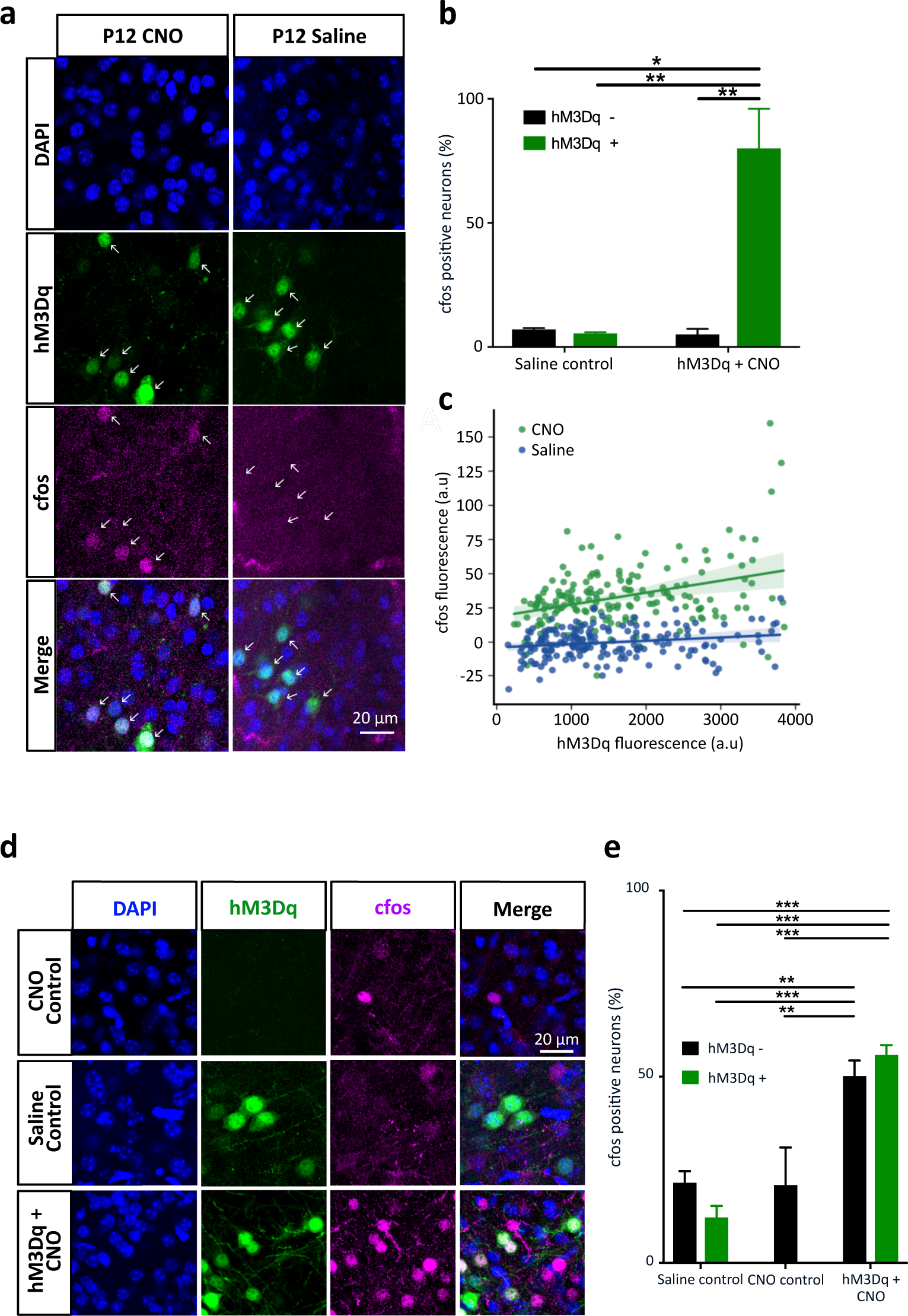
Cfos expression after CNO injection in hM3Dq+ and surrounding pyramidal cells. **(a)** Representative images of cfos expression in L2/3 cells in somatosensory cortex following a CNO or saline injection in P12 mice. **(b)** Percentage of cfos positive cells in saline and CNO injected animals (Chi-Square test, N = 300 cells, 3 mice, per condition). **(c)** Cfos fluorescence as a function of hM3Dq fluorescence. Linear regression, shade represents 95% confidence interval (Spearman’s correlation for CNO, p<0.05, r= 0.152). N = 57 cells CNO hM3Dq+, 543 cells CNO hM3Dq-network, 2 mice; 51 cells saline hM3Dq+, 549 cells saline hM3Dq-network, 2 mice). **(d)** Representative images of cfos expression in L2/3 cells following repeated injections of CNO/saline from P12-P18. **(e)** Percentage of cfos positive cells across conditions (Chi-Square test, N = 300 cells, 3 mice, per condition). *p<0.05, **p<0.01, ***p<0.001, bar plots show mean ± s.e.m.

**Extended Data Figure 4.**
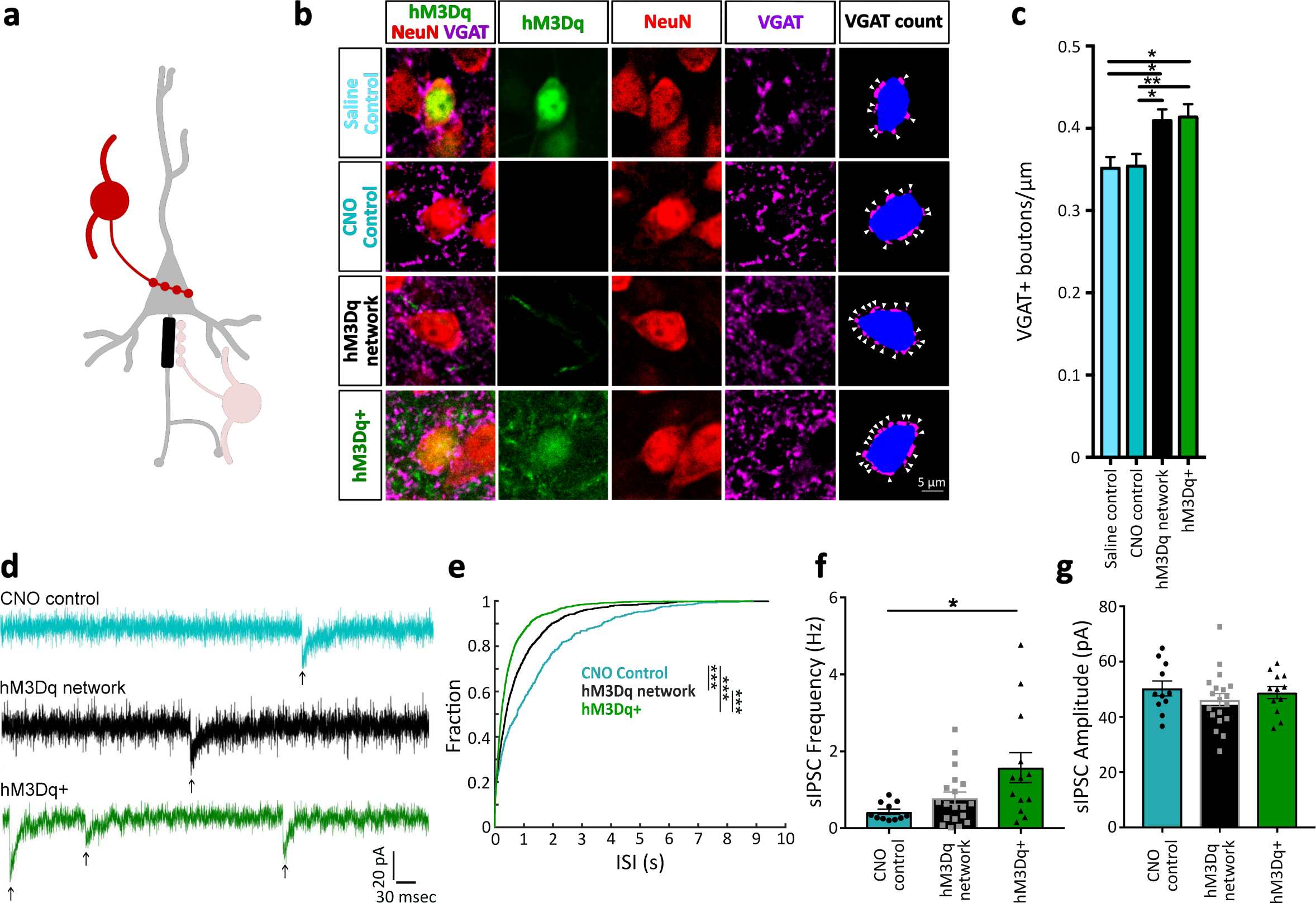
High network activity decreases the number of GABAergic synapses at the soma. **(a)** Basket cell synapses were quantified after chronic high network activity. **(b)** Confocal images of somas and synapses from L2/3 somatosensory cortex, from P18 mice. Cells were thresholded and automatically segmented as shown in ‘VGAT count’. **(c)** Density of synapses on the soma across groups. Values expressed as mean ± s.e.m. *p<0.05 **p<0.01 tested with one-way ANOVA with Sidak’s posthoc comparison test. CNO control: n = 30 neurons, 3 mice; Saline control: n = 20 neurons, 3 mice; hM3Dq: n = 30 neurons, 3 mice, hM3Dq-network: n = 30 neurons, 3 mice. **(d)** Representative traces of spontaneous IPSCs (sIPSCs). **(e)** Interspike interval (ISI), **(f)** average frequency and **(g)** amplitude of sIPSCs. (d-f) Values expressed as mean ± s.e.m. *p<0.05 ***p<0.001, tested with Kruskal-Wallis test followed by Dunn’s multiple comparisons test. CNO control: n = 11 neurons, 3 mice; hM3Dq-network: n = 19 neurons, 5 mice, hM3Dq+: n = 13 neurons, 3 mice.

**Extended Data Figure 5.**
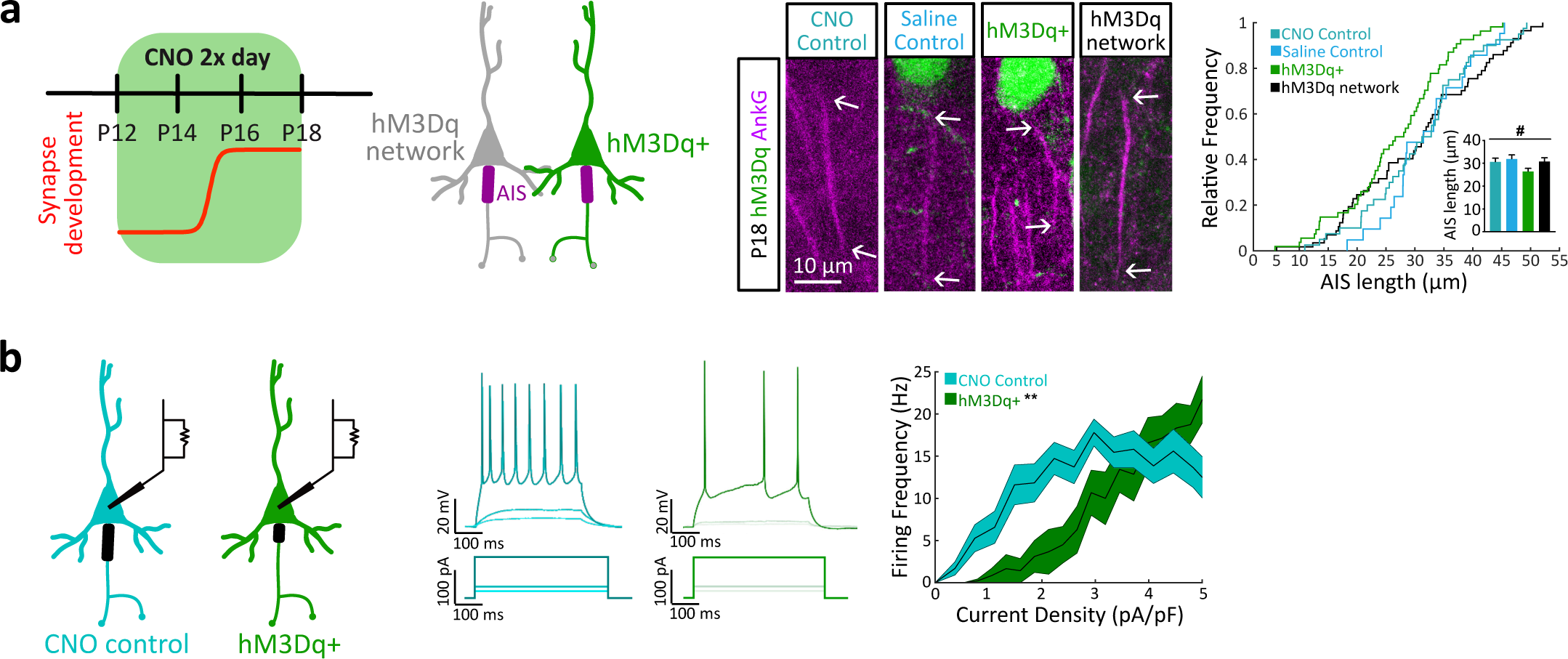
Increased network activity reduces AIS length. **(a)** Left, timeline for chronic activation of hM3Dq in L2/3 networks with a mixed population of hM3Dq+ (green) and hM3Dq-(grey) cells. Middle, representative images of AISs from L2/3 somatosensory cortex from P18 mice. White arrows delimit AIS. Right, length of the AIS across treatment groups. Values expressed as mean ± s.e.m., # denotes p<0.05 for one-way ANOVA without reaching significance in posthoc comparisons. CNO control: n = 57 neurons, 3 mice; saline control: n = 21 neurons, 4 mice; hM3Dq+: n = 54 neurons, 4 mice; hM3Dq-network: n = 40 neurons, 4 mice. **(b)** Left, CNO control (cyan) and hM3Dq+ neurons (green) were patched in whole-cell configuration to examine the intrinsic excitability properties after the treatment in (a). Middle, example firing profiles of a control (cyan) and hM3Dq+ (green) pyramidal cell to 500 ms current injections (steps shown: 10, 20 and 120 pA). Right, input-output curve for each group normalised to the neuron’s capacitance (current density). Centre line represents mean firing rate. Shaded area represents ± s.e.m. **p<0.05, repeated measures ANOVA. CNO control: n = 7 neurons, 2 mice; hM3Dq+: n = 8 neurons, 2 mice.

**Extended Data Figure 6.**
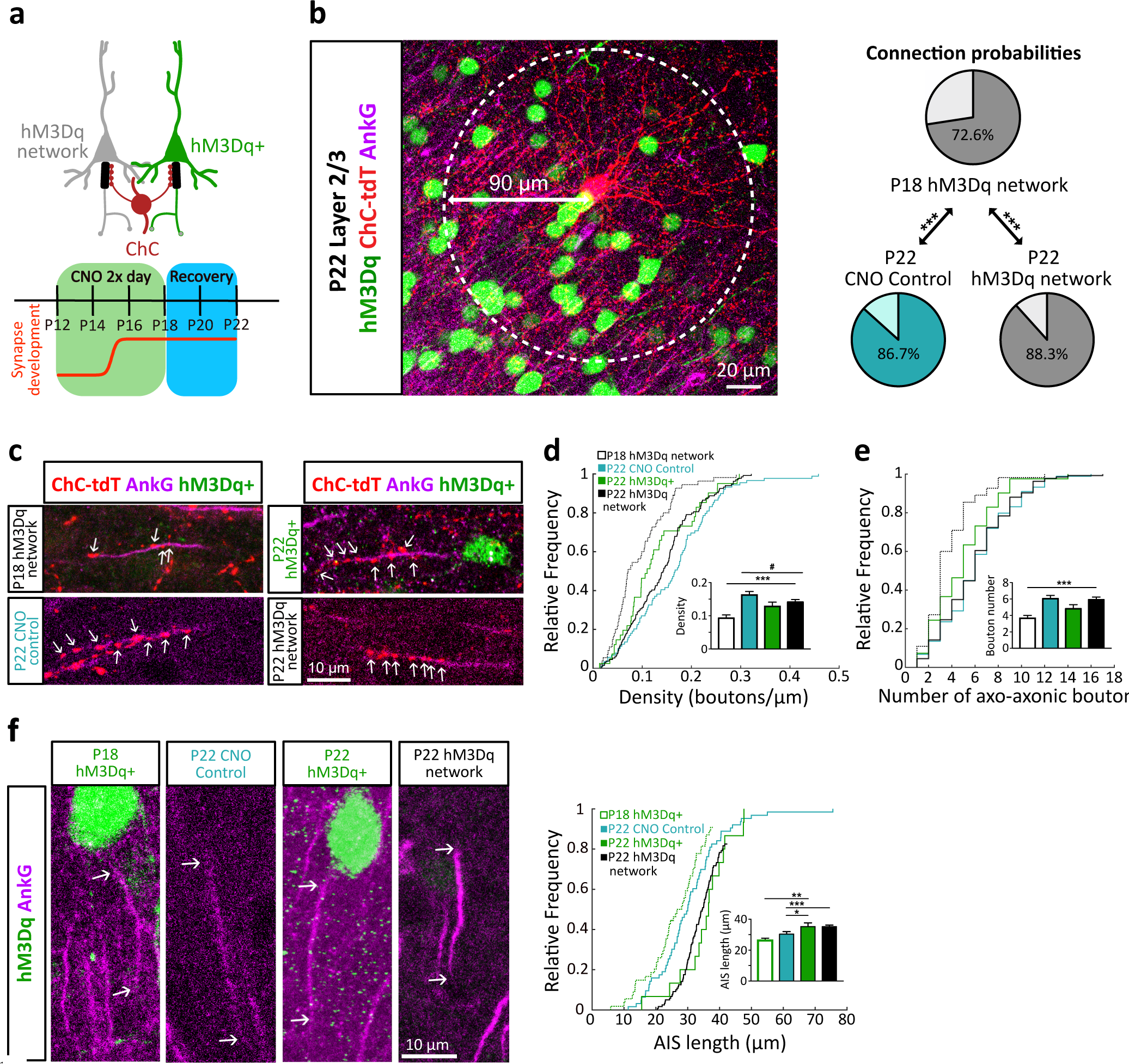
Axo-axonic plasticity is reversible. **(a)** Logic of experimental design and timeline of CNO application, including recovery period. **(B)** Example image of a ChC and connection probabilities within a 90 µm radius at P22, after the recovery period (Chi-square test, N = 180-390 AISs from 2-3 mice, per condition). **(c)** Example images of axo-axonic synapses and AISs following the recovery period. **(d,e)** Cumulative distribution of and average axo-axonic bouton density and number (# denotes p<0.05, Kruskal Wallis test with Dunn’s posthoc comparisons; ***p<0.01 for P18 versus P22 hM3Dq-network, Mann-Whitney test. N = 41-89 neurons, from 2-3 mice, per condition. **(f)** Left, example images of AISs from L2/3 somatosensory cortex at P22. Right, AIS length across conditions. * p<0.05, **p<0.01. ***p<0.001, P22 conditions tested with Kruskal-Wallis test followed by Dunn’s multiple comparisons test. P18 hM3Dq+ & P22 hM3Dq+ comparison tested with unpaired student’s t-test. Values expressed as mean ± s.e.m. CNO control: n = 63 neurons, 2 mice; hM3Dq+: n = 15 neurons, 3 mice, hM3Dq-network: n = 120 neurons, 3 mice.

**Extended Data Figure 7.**
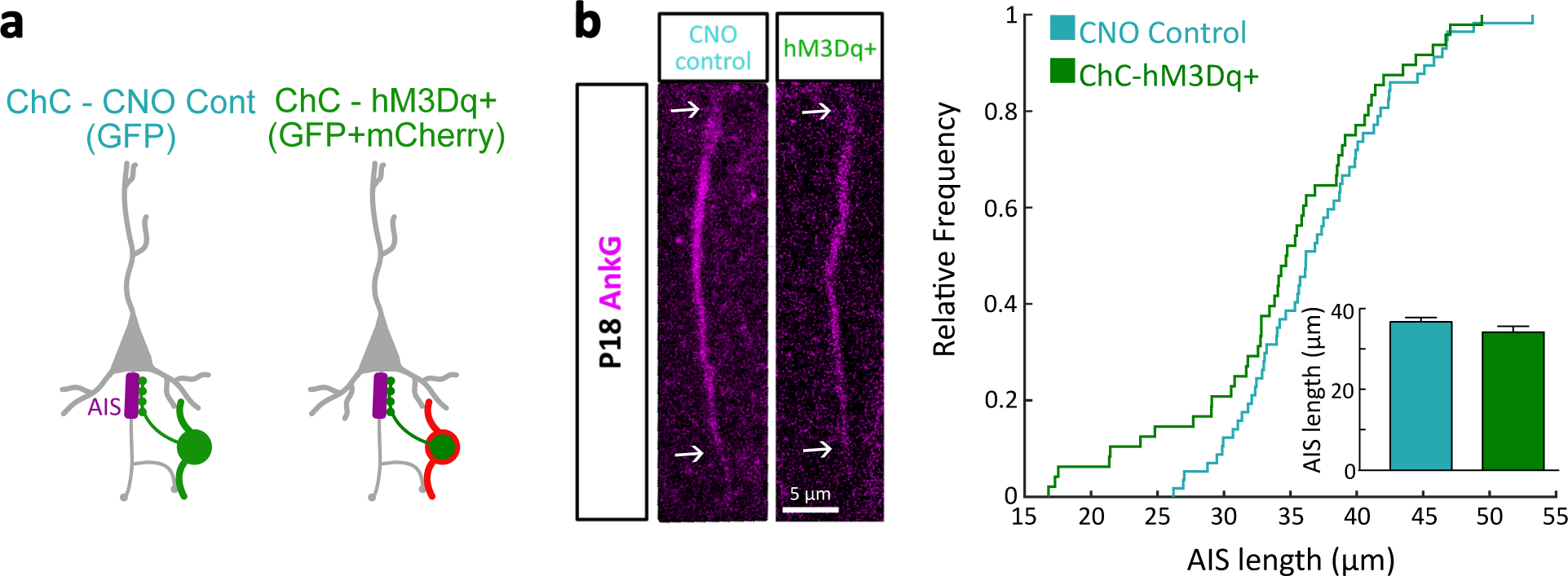
AIS plasticity requires direct pyramidal cell stimulation. **(a)** Experimental conditions generated by viral strategy, with ChCs expressing hM3Dq and GFP or control ChCs expressing GFP only. **(b)** Example images of AISs targeted by CNO control or hM3Dq+ ChCs. Right, AIS length across conditions. Differences tested with Mann-Whitney test. Values expressed as mean ± s.e.m. CNO control: n = 57 AIS, 3 mice; ChC-hM3Dq+ = 48 AIS, 3 mice.

**Extended Data Figure 8.**
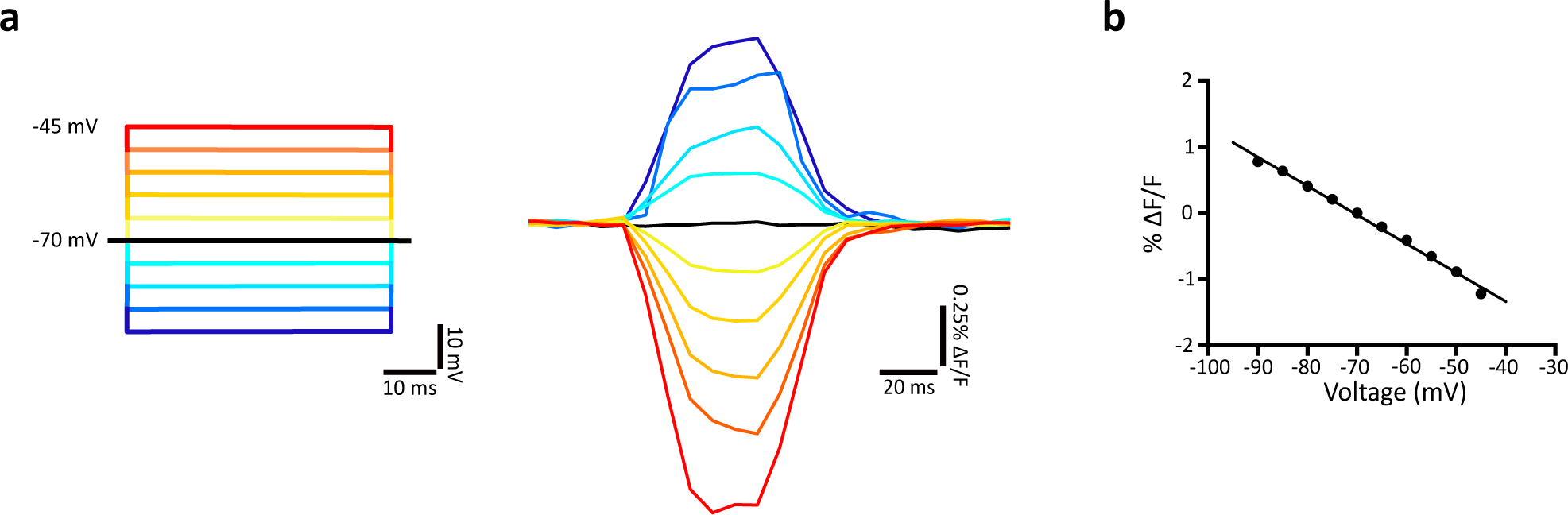
Ace2N-mNeon accurately reports small changes in voltage in acute slices. **(a)** Representative example of a fluorometric trace in response to voltage steps during a whole-cell patch-clamp experiment in a cell expressing Ace2N-mNeon. Yellow/red traces represent depolarizing voltage steps. Blue traces represent hyperpolarising voltage steps. **(b)** Maximum response (steady state) as a function of membrane voltage in the representative cell. Black line, linear regression (p<0.0001, r^2^ = 0.994).

**Extended Data Figure 9.**
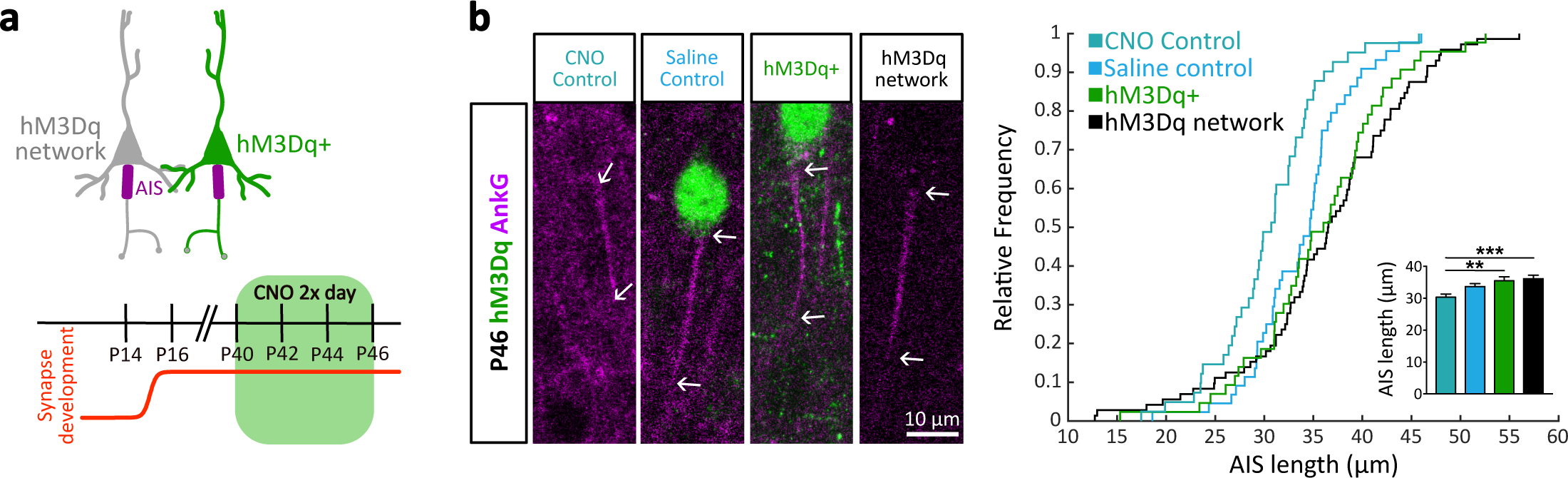
AIS plasticity does not occur in mature somatosensory cortex. **(a)** Timeline for activation of hM3Dq in L2/3 networks with a mixed population of hM3Dq+ (green) and hM3Dq-(grey) cells in adult mice. **(b)** Example images of AISs from L2/3 somatosensory cortex from P46 mice after adult treatment. Right, AIS length across conditions. **p<0.01, ***p<0.001 tested with Kruskal-Wallis and Dunn’s posthoc comparisons test. Values expressed as mean ± s.e.m. CNO control: n = 45 neurons, 3 mice; Saline control: n = 53 neurons, 5 mice, hM3Dq: n = 53 neurons, 3 mice, hM3Dq-network: n = 75 neurons, 3 mice.

## References

1 Isaacson, J. S. & Scanziani, M. How inhibition shapes cortical activity. Neuron 72, 231–243, doi:10.1016/j.neuron.2011.09.027 (2011).

2 Tremblay, R., Lee, S. & Rudy, B. GABAergic Interneurons in the Neocortex: From Cellular Properties to Circuits. Neuron 91, 260–292, doi:10.1016/j.neuron.2016.06.033 (2016).

3 Kubota, Y. Untangling GABAergic wiring in the cortical microcircuit. Current opinion in neurobiology 26, 7–14, doi:10.1016/j.conb.2013.10.003 (2014).

4 Inan, M. & Anderson, S. A. The chandelier cell, form and function. Current opinion in neurobiology 26, 142–148, doi:10.1016/j.conb.2014.01.009 (2014).

5 Wefelmeyer, W., Puhl, C. J. & Burrone, J. Homeostatic Plasticity of Subcellular Neuronal Structures: From Inputs to Outputs. Trends Neurosci 39, 656–667, doi:10.1016/j.tins.2016.08.004 (2016).

6 Turrigiano, G. Too many cooks? Intrinsic and synaptic homeostatic mechanisms in cortical circuit refinement. Annu Rev Neurosci 34, 89–103, doi:10.1146/annurev-neuro-060909-153238 (2011).

7 Kepecs, A. & Fishell, G. Interneuron cell types are fit to function. Nature 505, 318–326, doi:10.1038/nature12983 (2014).

8 De Marco Garcia, N. V., Karayannis, T. & Fishell, G. Neuronal activity is required for the development of specific cortical interneuron subtypes. Nature 472, 351–355, doi:10.1038/nature09865 (2011).

9 Karayannis, T., De Marco Garcia, N. V. & Fishell, G. J. Functional adaptation of cortical interneurons to attenuated activity is subtype-specific. Frontiers in neural circuits 6, 66, doi:10.3389/fncir.2012.00066 (2012).

10 Ben-Ari, Y., Gaiarsa, J. L., Tyzio, R. & Khazipov, R. GABA: a pioneer transmitter that excites immature neurons and generates primitive oscillations. Physiol Rev 87, 1215–1284, doi:10.1152/physrev.00017.2006 (2007).

11 Somogyi, P. A specific ‘axo-axonal’ interneuron in the visual cortex of the rat. Brain Res 136, 345–350 (1977).

12 Inan, M. et al. Dense and overlapping innervation of pyramidal neurons by chandelier cells. The Journal of neuroscience: the official journal of the Society for Neuroscience 33, 1907–1914, doi:10.1523/JNEUROSCI.4049-12.2013 (2013).

13 Fazzari, P. et al. Control of cortical GABA circuitry development by Nrg1 and ErbB4 signalling. Nature 464, 1376–1380, doi:10.1038/nature08928 (2010).

14 DeFelipe, J. Chandelier cells and epilepsy. Brain 122 (Pt 10), 1807–1822 (1999).

15 Marco, P. & DeFelipe, J. Altered synaptic circuitry in the human temporal neocortex removed from epileptic patients. Exp Brain Res 114, 1–10 (1997).

16 Taniguchi, H., Lu, J. & Huang, Z. J. The spatial and temporal origin of chandelier cells in mouse neocortex. Science 339, 70–74, doi:10.1126/science.1227622 (2013).

17 Urban, D. J. & Roth, B. L. DREADDs (designer receptors exclusively activated by designer drugs): chemogenetic tools with therapeutic utility. Annu Rev Pharmacol Toxicol 55, 399–417, doi:10.1146/annurev-pharmtox-010814-124803 (2015).

18 Kuba, H., Oichi, Y. & Ohmori, H. Presynaptic activity regulates Na(+) channel distribution at the axon initial segment. Nature 465, 1075–1078, doi:10.1038/nature09087 (2010).

19 Grubb, M. S. & Burrone, J. Activity-dependent relocation of the axon initial segment fine-tunes neuronal excitability. Nature 465, 1070–1074, doi:10.1038/nature09160 (2010).

20 Evans, M. D., Dumitrescu, A. S., Kruijssen, D. L. H., Taylor, S. E. & Grubb, M. S. Rapid Modulation of Axon Initial Segment Length Influences Repetitive Spike Firing. Cell Rep 13, 1233–1245, doi:10.1016/j.celrep.2015.09.066 (2015).

21 Wefelmeyer, W., Cattaert, D. & Burrone, J. Activity-dependent mismatch between axo-axonic synapses and the axon initial segment controls neuronal output. Proceedings of the National Academy of Sciences of the United States of America 112, 9757–9762, doi:10.1073/pnas.1502902112 (2015).

22 Muir, J. & Kittler, J. T. Plasticity of GABAA receptor diffusion dynamics at the axon initial segment. Front Cell Neurosci 8, 151, doi:10.3389/fncel.2014.00151 (2014).

23 Lu, J. et al. Selective inhibitory control of pyramidal neuron ensembles and cortical subnetworks by chandelier cells. Nature neuroscience 20, 1377–1383, doi:10.1038/nn.4624 (2017).

24 Turrigiano, G. G. & Nelson, S. B. Homeostatic plasticity in the developing nervous system. Nature reviews. Neuroscience 5, 97–107, doi:10.1038/nrn1327 (2004).

25 Rinetti-Vargas, G., Phamluong, K., Ron, D. & Bender, K. J. Periadolescent Maturation of GABAergic Hyperpolarization at the Axon Initial Segment. Cell Rep 20, 21–29, doi:10.1016/j.celrep.2017.06.030 (2017).

26 Gong, Y. et al. High-speed recording of neural spikes in awake mice and flies with a fluorescent voltage sensor. Science 350, 1361–1366, doi:10.1126/science.aab0810 (2015).

27 Woodruff, A. R. et al. State-dependent function of neocortical chandelier cells. The Journal of neuroscience: the official journal of the Society for Neuroscience 31, 17872–17886, doi:10.1523/JNEUROSCI.3894-11.2011 (2011).

28 Woodruff, A., Xu, Q., Anderson, S. A. & Yuste, R. Depolarizing effect of neocortical chandelier neurons. Frontiers in neural circuits 3, 15, doi:10.3389/neuro.04.015.2009 (2009).

29 Szabadics, J. et al. Excitatory effect of GABAergic axo-axonic cells in cortical microcircuits. Science 311, 233–235, doi:10.1126/science.1121325 (2006).

